# Cyt *b_6_/f* Complex Fine-Tunes PSI Stability and Photosynthetic Capacity Under Fluctuating Light

**DOI:** 10.1101/2025.02.16.638563

**Authors:** Masaru Kono, Hiromasa Kodama, Keiichiro Tanigawa, Ichiro Terashima, Wataru Yamori

## Abstract

Plants encounter dynamic light fluctuations in natural habitats, requiring robust mechanisms to protect Photosystem I (PSI) against photoinhibition. The cytochrome (Cyt) *b_6_/f* complex plays a pivotal role in regulating electron flow and ensuring PSI stability under fluctuating light. However, its precise role in balancing PSI protection and photosynthetic capacity remains unclear. In this study, transgenic tobacco (*Nicotiana tabacum* L. ‘W38’) with reduced Cyt *b_6_/f* content was used to examine PSI tolerance to photoinhibition and photosynthetic performance under fluctuating light. Notably, the reduced Cyt *b_6_/f* content conferred significant protection of PSI against photoinhibition induced by fluctuating light, albeit at the expense of the electron transport capacity. Remarkably, PSI stability was maintained even under extreme fluctuating light conditions in the plants with lowest Cyt *b_6_/f* contents, highlighting a trade-off between photoprotection and photosynthetic capacity. These findings reveal that a reduction in Cyt *b_6_/f* content enhances PSI protection while adjusting electron transport capacity, making plants more adaptable in environments dominated by low light and transient sunflecks. This study provides new insights into the adaptive strategies of plants under dynamic light environments and underscores the potential of targeting Cyt *b_6_/f* for enhancing crop tolerance to fluctuating light conditions.

## Introduction

Plants in natural habitats encounter dynamic and complex light environments. Open-field plants face gradual fluctuations caused by cloud movements and diurnal cycles, while understory plants endure rapid, short-term light fluctuations due to moving leaves and stems above them (Pearcy 1983, 1990, 1994; Chazdon 1988; Vierling and Wessman 2000). These changes, particularly in shaded environments, pose challenges to the photosynthetic apparatus, especially Photosystem I (PSI), which is vulnerable to photoinhibition under fluctuating light conditions (Suorsa et al. 2012; Kono et al. 2014, Kono and Terashima 2014). The photoinhibition of PSI results from imbalances in the electron flow upon the transition from the low-light phase to high-light phase, leading to over-reduction of the electron acceptors and the generation of reactive oxygen species (Sonoike and Terashima 1994; Terashima et al. 1994; Sonoike et al, 1995; Sonoike, 1995, 1996; Allahverdiyeva et al. 2005; Suorsa et al. 2012; Takagi et al. 2016).

Photosynthetic electron transport pathways include linear pathways and cyclic pathways. In linear electron pathway, electrons flow from Photosystem II (PSII) through the cytochrome *b_6_/f* (Cyt *b_6_/f*) complex to PSI. This process simultaneously generates a proton gradient across the thylakoid membrane, driving ATP synthesis via ATP synthase (Tikhonov 2013). However, fluctuating light has been shown to impose significant stress on PSI, leading to photoinhibition (Tsuyama et al. 2009, Sejima et al. 2014, Kono et al. 2014). This stress occurs when the balance between the PSI acceptor-side limitations and donor-side limitation is disrupted (Kono and Terashima 2016, Yamamoto and Shikanai 2019). Fluctuating light-induced PSI photoinhibition is reported to be in various plants such as *Arabidopsis thaliana*, rice, tobacco, and other field-grown species (Kono et al. 2017, Yamori et al. 2015, 2016a).

Maintaining the reaction center of PSI (P700) in the oxidized state (P700*^+^*) is crucial for mitigating fluctuating light-induced photodamage (Miyake 2020; Kono et al. 2022). Cyclic electron flow around PSI has been identified as a photoprotective mechanism under fluctuating light (Kono et al. 2014; Yamori et al. 2015, 2016a; Yamori and Shikanai 2016). Cyclic electron flow around PSI generates additional proton motive force (*pmf*), which enhances non-photochemical quenching (NPQ) (Miyake 2010; Strand et al.2016, 2017). NPQ safely dissipates excess excitation energy as heat, thereby preventing photodamage. Light-induced lumen acidification suppresses the linear electron transport at the Cyt *b_6_/f* complex, which is known as the ‘‘photosynthetic control.’’ Far-red light is another key factor in PSI stability and photoprotection (Kono et al. 2017). Far-red light maintains oxidized P700*^+^* upon the transitions from low-light phase to high light phase, mitigating acceptor-side limitations and enhancing NPQ, especially in sunfleck-type fluctuating light in deep shaded environments (Kono et al. 2017, 2020, 2022).

The Cyt *b_6_/f* complex plays a central role in regulating electron flow between PSII and PSI, as well as maintaining the transmembrane proton gradient (Yamori et al. 2011, Tikhonov 2013). This regulation is particularly crucial under fluctuating light conditions, where PSI is vulnerable to photoinhibition. Reducing Cyt *b_6_/f* content has been suggested as a strategy to enhance PSI protection under fluctuating light by limiting excessive electron flow to PSI, thereby mitigating photodamage. However, this reduction comes at the cost of decreased photosynthetic rates, highlighting an inherent trade-off between photoprotection and photosynthetic capacity. Thus, the ability to finely tune Cyt *b_6_/f* abundance represents a key adaptive strategy for optimizing the balance between light utilization and photoprotection in dynamic light environments. While the role of Cyt *b_6_/f* in the photosynthetic control is well established, its precise contributions to PSI photoprotection under fluctuating light remain insufficiently understood and warrant further investigation.

Despite the growing understanding of PSI protection mechanisms, several knowledge gaps persist. These include the precise role of the Cyt *b_6_/f* complex in balancing PSI protection and photosynthetic capacity, as well as the effects of mechanisms like FR-induced P700*^+^* oxidation and NPQ enhancement. Additionally, the adaptive strategies employed by shade-grown plants, which operate with inherently low photosynthetic capacities, warrant further investigation. In this study, we investigate the role of the Cyt *b_6_/f* complex in balancing PSI photoprotection and photosynthetic capacity under FL conditions. We used transgenic tobacco (*Nicotiana tabacum*) with Cyt *b_6_/f* complex content reduced to approximately 30% and 60% of the wild-type level (Yamori et al. 2011), and assessed the tolerance to PSI photoinhibition and the photosynthetic capacity. This research provides insights into the adaptive mechanisms of plants in dynamic light environments, emphasizing the balance of Cyt *b_6_/f* level and NPQ in optimizing photoprotection and photosynthetic capacity.

## Materials and Methods

### Plant Materials and Growth Conditions

Tobacco (*Nicotiana tabacum* L. cv. W38) plants and several transformants of anti-Rieske FeS tobacco that have reduced amounts of the Cyt *b_6_/f* were grown in growth cabinets (Yamori et al. 2011). Plants were grown in sterilized soil under controlled environmental conditions: an 8-hour photoperiod with a photosynthetic photon flux density (PPFD) of 200 µmol photons m⁻² s⁻¹, air temperatures of 23°C, and a relative humidity of 60%. Plants were grown in 5-L pots in a 1:1 ratio of Metro-Mix (Hyponex Japan, Osaka, Japan) and Vermiculite GL (Nittai Co., Ltd., Osaka, Japan), and were irrigated two to three times weekly and were fertilized with Hyponex 6-10-5 solution (Hyponex Japan, Osaka, Japan) diluted to the 1: 1000 strength every irrigation from two weeks after germination. For experiments, fully expanded leaves (third to fifth leaves from the apex) were selected to standardize measurements across replicates.

### Analysis of photosynthetic components

Immediately after photosynthetic measurements, leaf discs were taken and immersed in liquid nitrogen and stored at −80°C. The frozen leaf sample was ground in liquid nitrogen and homogenized in an extraction buffer (Yamori et al. 2011). The contents of Rieske FeS of Cyt *b_6_/f* complex and δ-subunit of ATP synthase were quantified by immunoblotting with anti-Rieske FeS antibody and anti-ATP synthase (δ) antibody (Agrisera). Chlorophylls were extracted in 80% (v/v) acetone and determined (Porra et al., 1989). The extract of one wild-type leaf was selected as a standard (100%) and included as a dilution series on gels. The protein content of other samples was referenced against this standard.

### Fluctuating Light Treatment

Leaves were exposed to fluctuating light to investigate PSI photoinhibition and photosynthetic responses (Tsuyama et al. 2009, Sejima et al. 2014, Kono et al. 2017). For measurement of PSI photoinhibition, fluctuating light consisted of alternating high light (2000, 3000, or 20,000 µmol photons m⁻² s⁻¹) and low light (30 µmol photons m⁻² s⁻¹) phases at intervals of 800 ms/10 s for a total duration of 120 min. For these light treatments, red light-emitting diodes (LEDs; Excelitas/ELCOS GmbH, Germany) with the wavelength peak at 635 nm were used.

For measurements of photosynthetic responses, we used the fluctuating light, in which high light at 800 μmol m^−2^ s^−1^ for 10 min and low light at 30 μmol m^−2^ s^−1^ for 15 min alternated.

### Chlorophyll Fluorescence and 830 nm Absorbance Change Measurements

Chlorophyll fluorescence and absorption changes at 830 nm were measured simultaneously in intact leaves using a DUAL-PAM-100 (Walz, Effeltrich, Germany) in the ventilated room air. Saturation pulses (SPs) from red LEDs (7000 μmol m^−2^ s^−1^, 500 ms duration) were applied to determine the maximum chlorophyll fluorescence with closed PSII centers after dark treatment (F*_m_*) and during illumination (F*_m_*’). The maximum photochemical quantum yield of PSII (F*_v_* /F*_m_*) in the dark and the effective quantum yield of PSII (Y(II)) in the actinic light were calculated as (F*_m_* − F*_o_*)/F*_m_* and (F*_m_*’ − F*_s_*’)/F*_m_*’ (Kitajima and Butler 1975; Genty et al. 1989), respectively, where F*_o_* is the minimal chlorophyll fluorescence in the dark and F*_s_*’ is the steady-state chlorophyll fluorescence level in the actinic light from red LEDs. Non-photochemical quenching (NPQ) was calculated as (F*_m_* – F*_m_*’)/F*_m_*’ (Schreiber and Bilger 1987; Bilger and Bjorkman 1990). The coefficient of photochemical quenching (qL), a measure of the fraction of open PSII reaction centers, based on the “lake model” of PSII antenna pigment organization, was calculated as (F*_m_*’ – F*_s_*’)/(F*_m_*’ – F*_o_*’) • F*_o_*’/F*_s_*’ (Kramer et al. 2004). F*_o_*’ is the minimal fluorescence yield in the actinic light and was estimated using the formula of Oxborough and Baker (1997) as F*_o_*/(F*_v_* /F*_m_* + F*_o_*/F*_m_*’).

With the Dual-PAM-100, P700*^+^* was monitored as the absorption difference between 830 and 875 nm in a transmission mode. In analogy to the quantum yields of PSII, the quantum yields of PSI were determined using the saturation pulse method (Klughammer and Schreiber 1994). The maximum oxidizable P700, P*_m_*, was determined by application of the SP in the presence of far-red light at 25.6 W m^−2^ with the wavelength peak at 740 nm. The decrease in P*_m_* is an indicator of PSI photoinhibition. Y(I), Y(ND) and Y(NA) are determined in the light. These add up to unity with the photochemical quantum yield (i.e. Y(I) + Y(ND) + Y(NA) = 1).

### Measurement of the Electrochromic Absorbance Shift

Carotenoid electrochromic absorbance shift (ECS or P515) measurements were performed with the DUAL-PAM-100 using the P515/535 module, to assess the proton motive force (*pmf*) and its components (ΔΨ and ΔpH). ECS signals were recorded at 515 nm with reference wavelengths at 550 nm (Junge and Witt 1968, Klughammer et al. 2013). For determinations of the size of the *pmf* (Sacksteder et al. 2000, Cruz et al. 2001), the dark-interval relaxation kinetics (DIRK) transient-analysis was conducted (for detail, see Baker et al. 2007). The *pmf* amplitude was determined from ECS decay kinetics (Sacksteder et al. 2000; Cruz et al. 2001). Contributions of the ΔΨ (electric potential) and ΔpH (proton gradient) were inferred by analyzing the relaxation of the ECS signal during dark intervals. ECS signals were normalized by a single-turnover flash for 8 µs.

### Statistical Analysis

All data are expressed as means ± standard error (SE). Statistical analyses were conducted using R software version 4.1.2. One-way ANOVA was performed to compare differences between WT and transgenic plants, followed by Tukey’s post-hoc test to identify significant pairwise differences. A *p*-value of < 0.05 was considered statistically significant for all analyses.

## Results

### Characterization of Cyt *b*_6_*/f* Complex Content in Transgenic Plants

To investigate the role of the Cyt *b_6_/f* complex in photosynthesis, transgenic tobacco plants with reduced levels of the Rieske FeS protein, a key subunit of Cyt *b_6_/f*, were generated (**Figure 1A**). Three groups were classified with respect to Rieske FeS content: the wild type (WT; 100%), plants with intermediate Rieske FeS levels (43.6 to 76.2%), and plants with low Rieske FeS levels (18.5 to 42.2%) (**Figure 1B**). On the other hand, the contents of several photosynthetic components, including δ-subunit of ATP synthase, Rubisco and chlorophylls, were similar among the wild type and transgenic plants (**Figure 1C, D, E**). We therefore assume that alterations in photosynthetic properties are primarily the result of the reduction in Cyt *b_6_/f* complex in transgenic plants. These results validate the transgenic lines as an effective model for studying the role of Cyt *b_6_/f* in photosynthetic regulation under varying light conditions.

**Figure 1.**
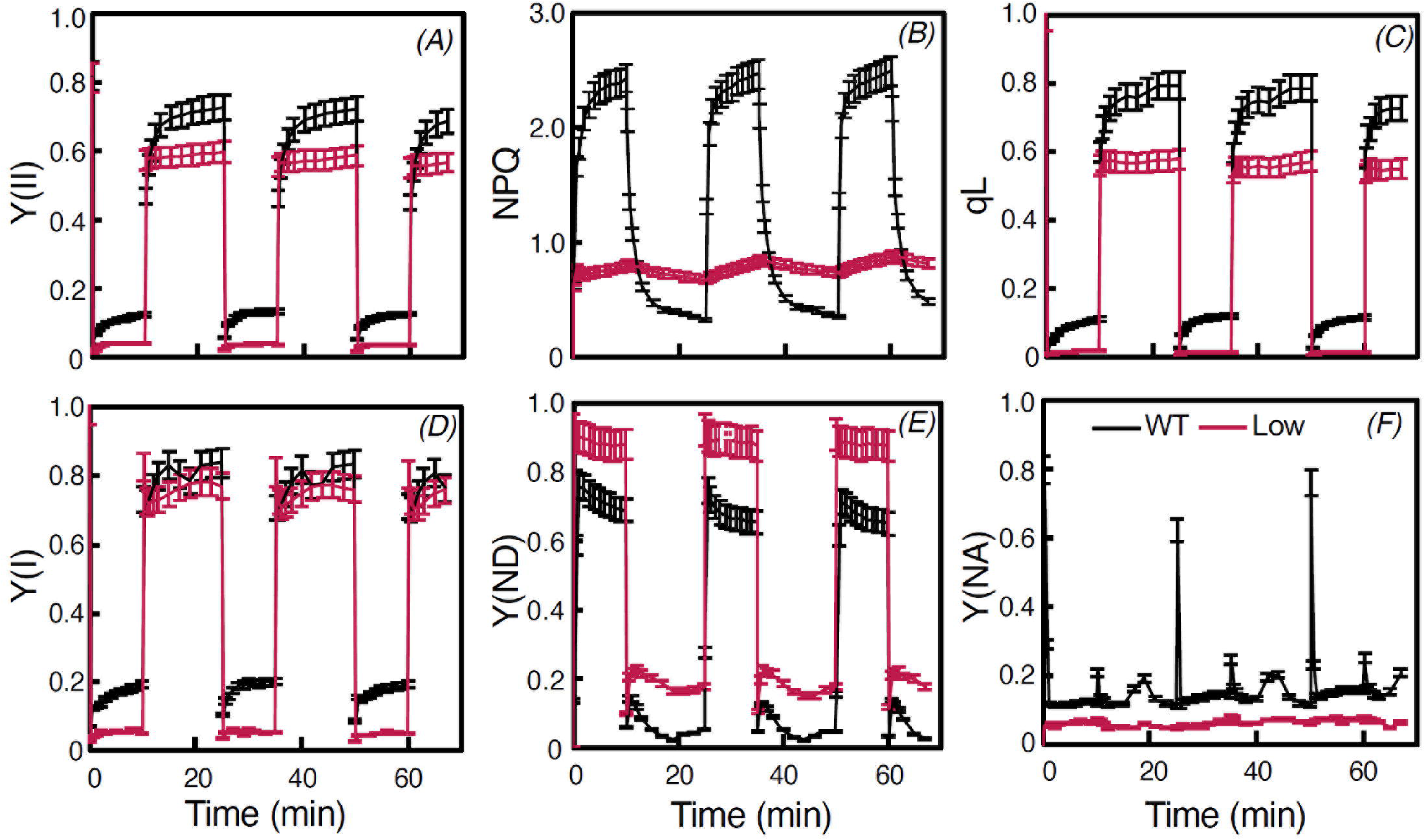
Characterization of the Cyt *b*_6_*/f* Complex Content in Transgenic Plants. (A) Proteins in the leaf extract were separated by sodium dodecyl sulphate–polyacrylamide gel electrophoresis (SDS-PAGE) and immunodetected with specific antibodies. The extracted proteins for each sample we reloaded on gel using an equal leaf area basis; the series of dilutions for wild-type plants are indicated. (B) Relative content of the Rieske FeS protein of Cyt *b_6_/f* complex, (C) Relative content of the δ-subunit of ATP synthase, (D) Rubisco content (µmol m⁻²) and (E) Chlorophyll *a* + *b* content (mmol m⁻²) in wild-type (WT) and transgenic plants categorized into “Middle” and “Low” groups based on Cyt *b_6_/f* content: the wild type (WT; approximately 100%), plants with intermediate Rieske FeS level (43.6 to 76.2%), and plants with low Rieske FeS level (18.5 to 42.2%). Data are presented as means ± SE (*n* = 3 ∼ 5, \**p* < 0.05).

### Proton Motive Force ( *pmf*) and Its Components

The proton motive force (*pmf*) and its components, ΔpH and ΔΨ, were measured to assess the impact of Cyt *b_6_/f* reduction on energy conversion (**Figures 2A-C**, for detailed statistics analyses, see **Figures S1A-C**). In WT plants, the highest *pmf* was observed at all PPFDs, followed by those of “Middle” and “Low” groups (**Figures 2A and S1A**). The ΔpH and ΔΨ components also decreased proportionally with the Cyt *b_6_/f* reduction. Notably, ΔΨ was nearly absent in “Low” plants, making *pmf* in this group almost entirely ΔpH (**Figures 2C and S1C**). These findings indicate that reductions in Cyt *b_6_/f* content impair energy transduction efficiency, particularly by diminishing the ΔΨ, component of *pmf*.

**Figure 2.**
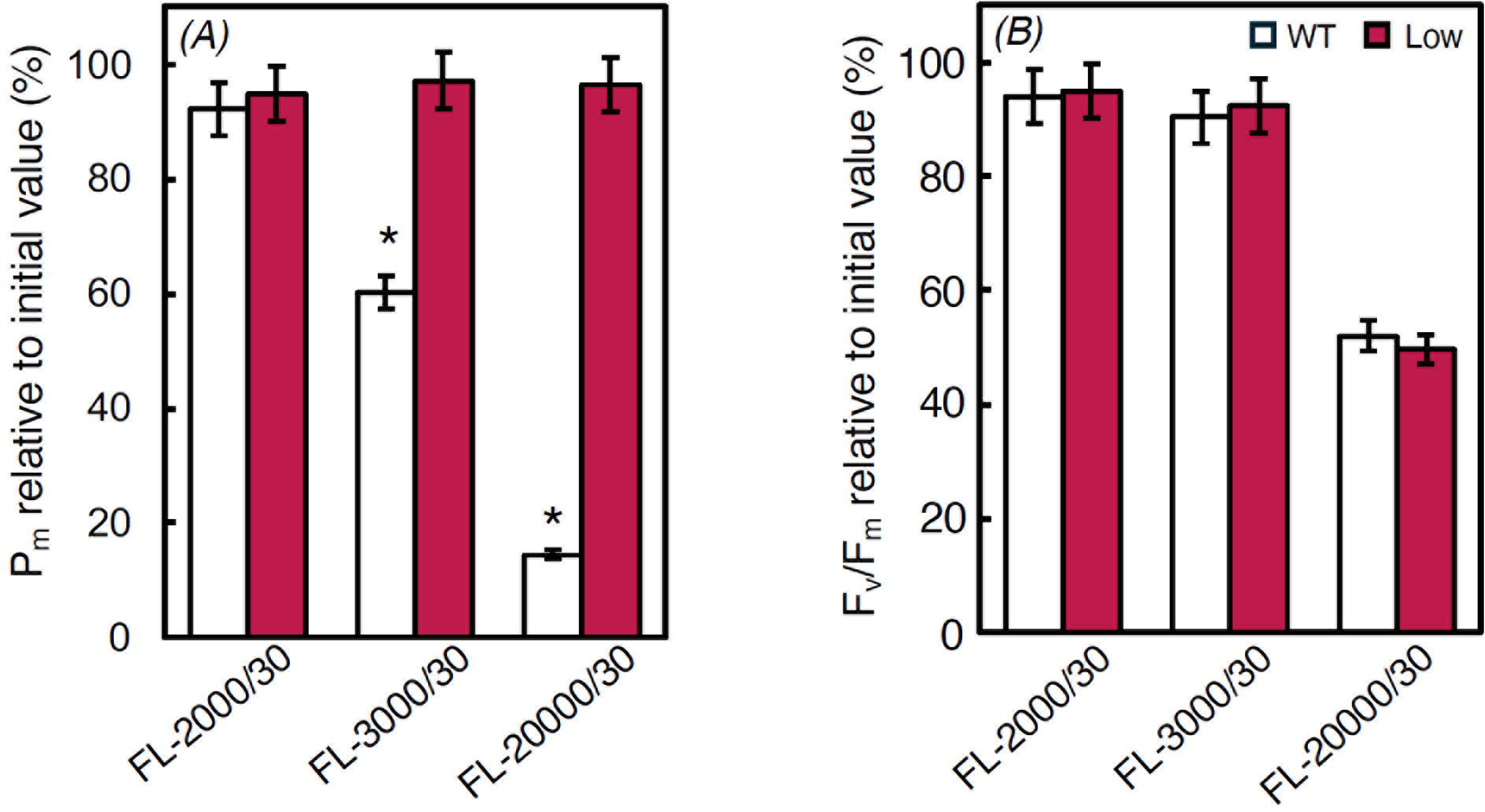
Proton Motive Force (*pmf*) and Its Components in WT and Transgenic Plants. Light response curves of (A) *pmf*, (B) ΔpH, and (C) ΔΨ components in WT, “Middle,” and “Low” plants. Each parameter was normalized by a single-turnover flash for 8 μs, with measurements conducted at varying light intensities (µmol m⁻² s⁻¹). Data are presented as means ± SE (*n* = 3).

### Light Response Curves for PSII and PSI Parameters

The light response curves for PSII and PSI parameters revealed the extent to which Cyt *b_6_/f* reductions affect photosynthetic electron transport (**Figure 3**). For PSII, Y(II) (**Figures 3A and S2A**) and qL (**Figures 3C and S2C**) were stable in “Middle” plants but significantly reduced in “Low” plants, indicating impaired electron transport. NPQ (**Figures 3B and S2B**) was notably reduced even in “Middle” plants, with levels halving compared to WT and declining further in “Low” plants, suggesting reduced NPQ (**Figures 3B and S2B**) was notably reduced in “Middle” plants, about half the WT. NPQ was further lowered in “Low” plants.

**Figure 3.**
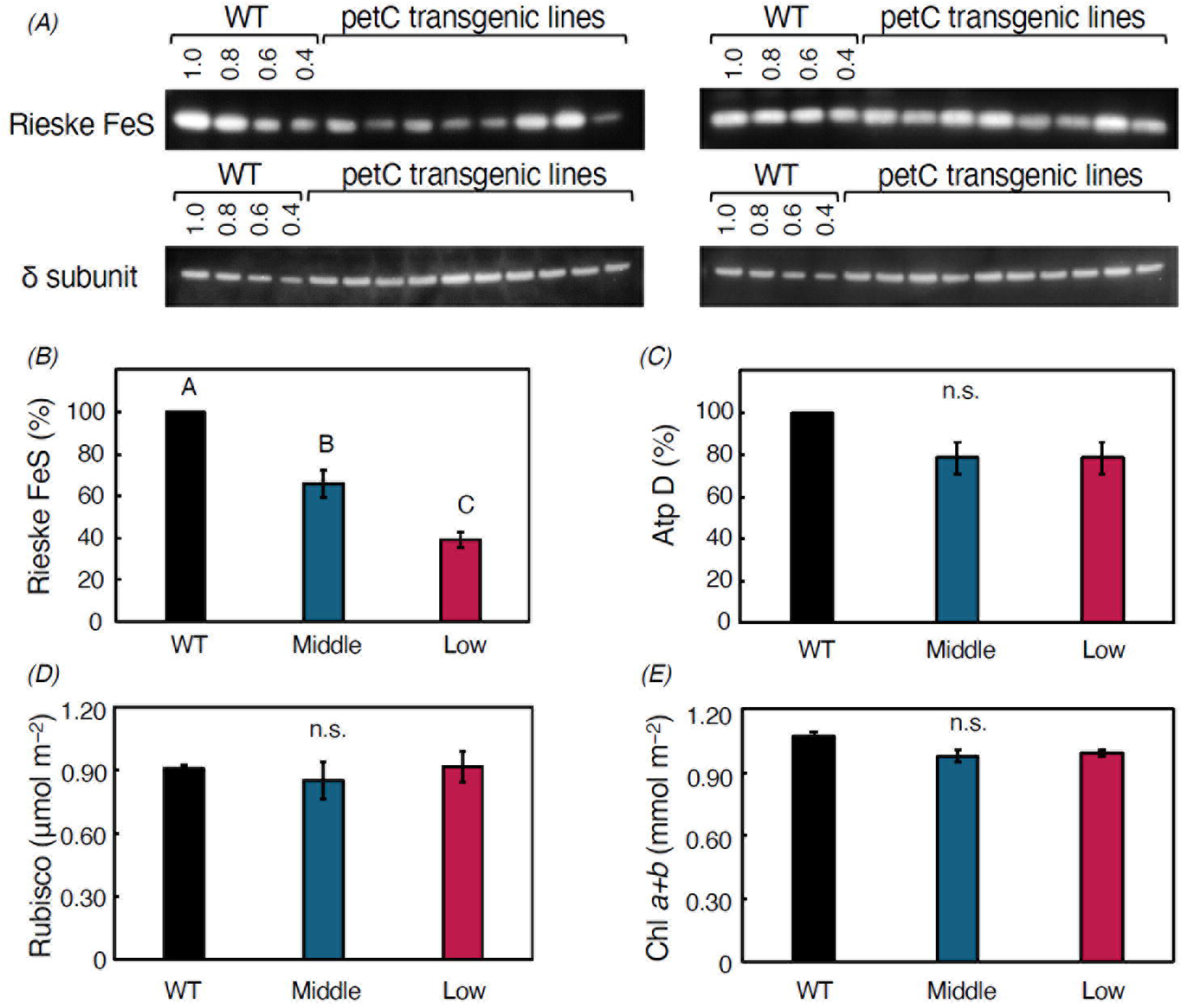
Light Response Curves of PSII and PSI Parameters. PSII parameters: (A) quantum yield of PSII (Y(II)), (B) non-photochemical quenching (NPQ), and (C) the redox state of the plastoquinone pool (qL) in WT, “Middle,” and “Low” plants. PSI parameters: (D) quantum yield of PSI (Y(I)), (E) donor-side limitation of PSI (Y(ND)), and (F) acceptor-side limitation of PSI (Y(NA)). Data are presented as means ± SE (*n* = 3).

Y(I) in “Middle” plants were similar to those in WT, but those in “Low” plants were decreased severly (**Figures 3D and S2D**). In the “Low” group, Y(ND) increased substantially (**Figures 3E and S2E**), while Y(NA) declined (**Figures 3F and S2F**), indicating severe donor-side limitations on PSI electron flow. These results demonstrate that moderate reductions in Cyt *b_6_/f* have minimal effects on electron flows in PSII and PSI, while severe reductions suppressed both PSII and PSI electron transport rates and NPQ.

### Photosynthetic Performance Under Fluctuating Light

PSII and PSI performance in dark-treated leaves were evaluated under the fluctuating light conditions, in which high and low light phases alternate (**Figure 4**). Y(II) and qL in “Low” plants reached lower steady-state levels during the high light phase and, in the low light phase, these were also lower in “Low” plants than in WT (**Figure 4A and C**). NPQ induction during high light was delayed and lower in “Low” plants, and its relaxation during low light was small (**Figure 4B**). These patterns indicate that “Low” plants had a lower capacity for the electron transport and PSII photoprotection under dynamic light conditions. For PSI, the patterns observed under fluctuating light were consistent with those under constant light (**Figures 4**). These findings confirm that Cyt *b_6_/f* plays a critical role in regulating PSII performance under fluctuating light.

**Figure 4.**
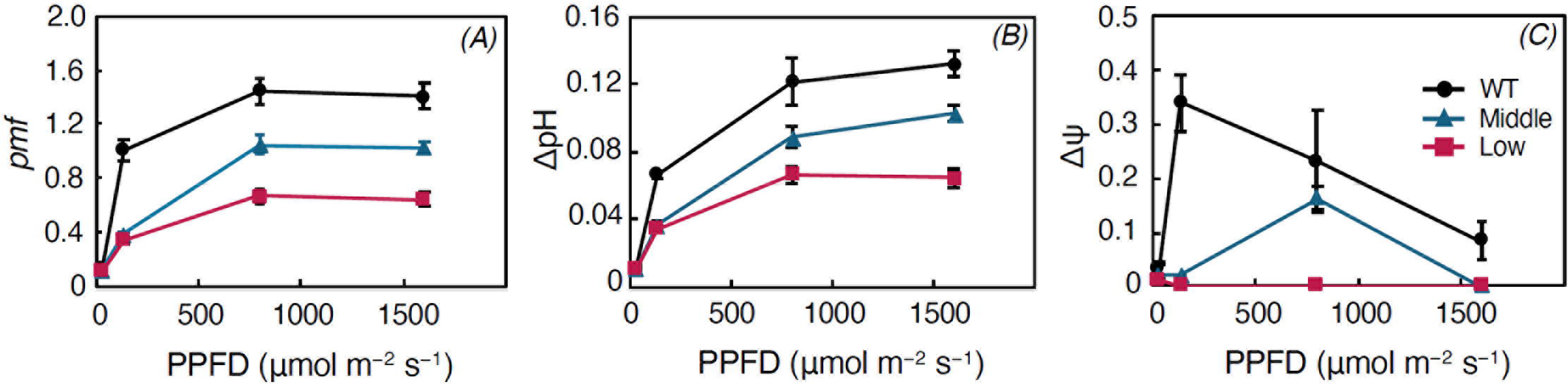
Photosynthetic Responses to Fluctuating Light in WT and “Low” Plants. (A-C) PSII parameters: Y(II), NPQ, and qL under alternating high (800 µmol m⁻² s⁻¹, 10 min) and low (30 µmol m⁻² s⁻¹, 15 min) light phases in WT, “Middle,” and “Low” plants. (D-F) PSI parameters: Y(I), Y(ND), and Y(NA) under the same fluctuating light conditions in WT, “Middle,” and “Low” plants. Data are presented as means ± SE (*n* = 3).

### PSI Photoinhibition Under Fluctuating Light

To assess photoinhibition, dark-treated leaves were exposed to fluctuating light with high light pulses superimposed on a weak background light. PSI photoinhibition was evaluated by measuring P*_m_*, and PSII photoinhibition was assessed through F*_v_* /F*_m_* (**Figure 5**). For PSI, WT plants showed no photoinhibition at 2,000 μmol m⁻² s⁻¹ but showed significant reductions in P*_m_* at 3,000 μmol m⁻² s⁻¹. At 20,000 μmol m⁻² s⁻¹, P*_m_* in WT plants declined to less than 20% of initial levels. Remarkably, “Low” plants exhibited complete resistance to PSI photoinhibition, maintaining stable P*_m_* levels even at the highest light intensity (**Figure 5A**).

**Figure 5.**
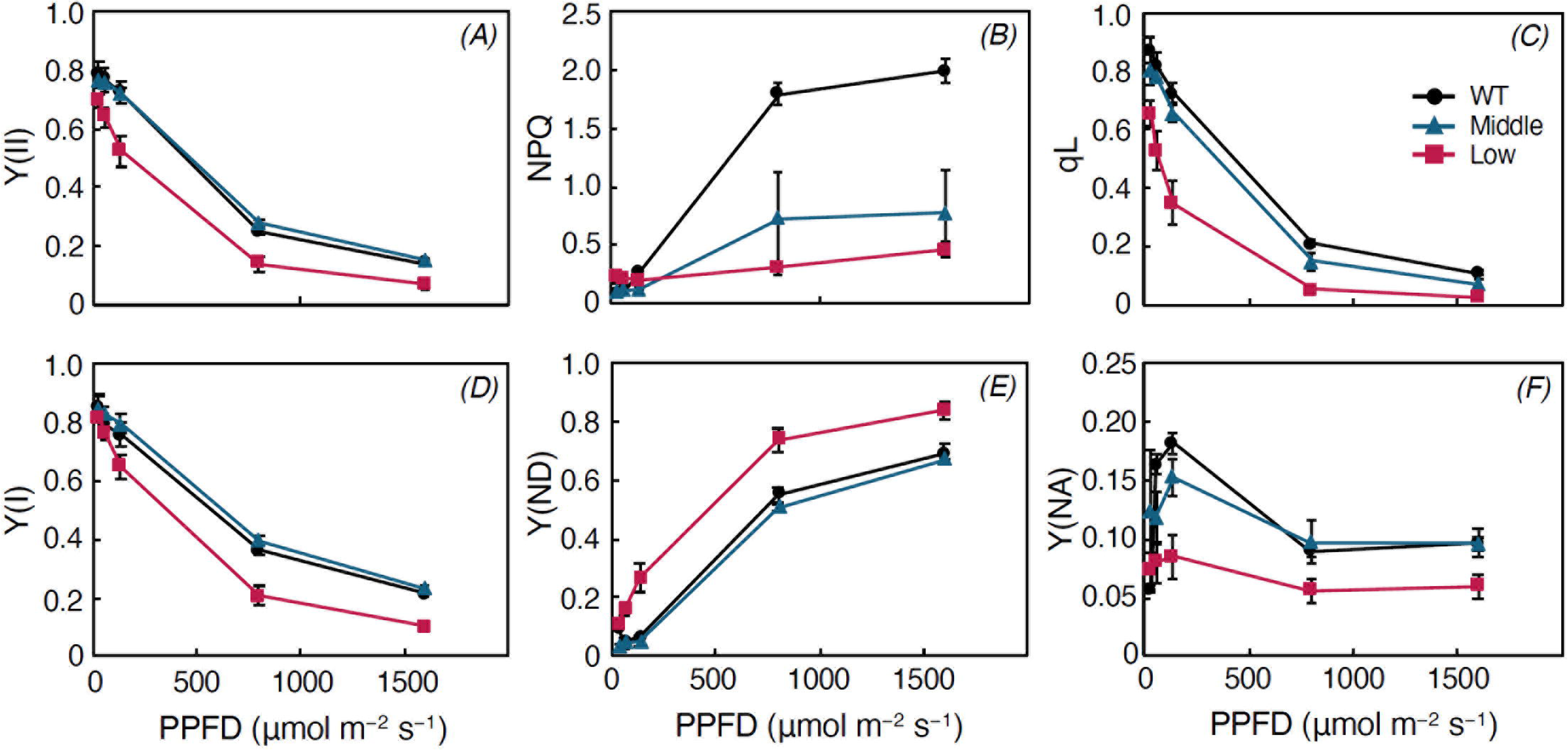
PSI and PSII Photoinhibition Under Sunfleck-Like Fluctuating Light. (A) Relative P*_m_* values (%) and (B) Relative F*_v_* /F*_m_* values (%) before and after exposure to fluctuating light with strong light pulses (800 ms) at 2,000, 3,000, or 20,000 µmol m⁻² s⁻¹, in WT and plants with low Rieske FeS level. Data are presented as means ± SE (*n* = 3, \**p* < 0.05).

For PSII, F*_v_* /F*_m_* values remained stable at 2,000 and 3,000 μmol m⁻² s⁻¹ but decreased to ∼50% in both WT and “Low” plants at 20,000 μmol m⁻² s⁻¹, indicating comparable susceptibility to photoinhibition (**Figure 5B**). These results highlight a selective role of Cyt *b_6_/f* in protecting PSI from photoinhibition under extreme light conditions, while PSII vulnerability appears less dependent on Cyt *b_6_/f* content against the flash-type fluctuating light.

## Discussion

This study explored the critical role of the Cyt *b_6_/f* complex in photosynthetic regulation, emphasizing its contributions to PSI stability under fluctuating light conditions. By examining transgenic tobacco plants with varying Cyt *b_6_/f* contents, we uncovered novel insights into how this complex mediates the photosynthetic capacity, PSII photoprotection and PSI photoprotection. Previous works demonstrated that Cyt *b_6_*/*f* plays pivotal roles in maintaining steady-state electron transport and CO₂ assimilation in C*_3_* plants (Yamori et al., 2011, 2016b). Consistent with these findings, we confirmed that Cyt *b_6_/f* content is a key determinant of electron transport capasity, as evidenced by reduced Y(II) and qL in “Low” plants (**Figures 3A, 3C, S2A and S2C**). While the previous works were made in constant light (Yamori et al., 2011, 2016b), the present study addressed the challenges posed by fluctuating light. We demonstrated that Cyt *b_6_/f* reductions modify the dynamic regulation of NPQ and the balance between PSI and PSII during rapid light transitions (**Figure 4**). These findings highlight the essential role of Cyt *b_6_/f* in facilitating rapid photoprotective adjustments to fluctuating light environments. Furthermore, our study revealed that reductions in Cyt *b_6_/f* content have divergent effects on PSII and PSI photoinhibition (**Figure 5**). Specifically, “Low” plants demonstrated exceptional resistance to PSI photoinhibition, even under extreme fluctuating light conditions. These results advance the understanding of Cyt *b_6_/f* as a dynamic regulator of photosynthesis and photoprotection, offering new insights into its potential for enhancing crop resilience in variable light environments.

### The Essential Role of Cyt *b*_6_*/f* in Photosynthetic Electron Transport

Our results provide new insights into the role of the Cyt *b_6_/f* complex in balancing electron transport capacity and photoprotection under fluctuating light conditions. Our findings align with Tikhonov’s (2013) review, which emphasizes the central role of Cyt *b_6_/f* in regulating photosynthetic electron flow. In the present study, reductions in Cyt *b_6_/f* content modified the *pmf*, by severely diminishing its ΔΨ component, resulting from slower electron flow between PSII and PSI. This confirms that Cyt *b_6_/f* is a key determinant of ΔΨ, consistent with its established role in facilitating proton transfer across the thylakoid membrane (Tikhonov 2014). Tikhonov (2013, 2014) described the role of PQH*_2_* oxidation at the Q*_o_* site of Cyt *b_6_/f* as a rate-limiting step that modulates the intersystem electron transport chain, particularly during high-light conditions. This regulation ensures that electron flow remains proportional to the ATP and NADPH production required for carbon fixation and metabolic processes while preventing over-reduction of PSI acceptors. Our results in “Low” plants, where Cyt *b_6_/f* content was severely reduced, revealed slower electron transport capacities and greater donor-side limitations on the PSI electron flow (**Figure 5**), corroborating the critical role of Cyt *b_6_/f* in maintaining the coordination between the electron supply from PSII and the downstream electron flows.

The differential effects of Cyt *b_6_/f* reduction on PSII and PSI activities further highlight its centrality in regulating electron transport. While “Low” plants exhibited significant impairments in both PSII and PSI performance, PSI in the “Middle” plants remained stable (**Figures 3 ans S2**), indicating that PSI electron transport is more resistant to moderate reductions in Cyt *b_6_/f*. This supports previous suggestions that PSI transport is less dependent on Cyt *b_6_/f* -mediated electron flow under steady-state conditions (Yamori et al. 2011, 2016b, Kono and Terashima 2016, Kono et al. 2020).

### The Role of Cyt *b*_6_*/f* in Balancing Energy Dissipation and PSI Stabilization

The adaptive significance of Cyt *b_6_/f* -mediated feedback control under variable environmental conditions has been emphasized (Yamori et al. 2011, 2016b; Tikhonov 2014). Our findings extend this understanding by showing that reduced Cyt *b_6_/f* content weakens the plant’s ability to adapt to fluctuating light environments. While “Low” plants displayed low NPQ levels (**Figure 4**), they demonstrated remarkable resistance to PSI photoinhibition, even under extreme light fluctuations (**Figure 5**). This indicates a trade-off facilitated by Cyt *b_6_/f*, where its modulation shifts the balance between energy dissipation and PSI stabilization. Such a trade-off may be adaptive in environments where PSI stability is prioritized, such as in shaded habitats with transient sunflecks (see “Resistance to PSI Photoinhibition: A Novel Trade-Off”).

Our results demonstrate that Cyt *b_6_/f* plays a pivotal role in PSII photoprotection under fluctuating light conditions (**Figure 5**). In “Low” plants, the delayed induction of NPQ and lower regulation of Y(II) and qL suggested an impaired ability to dissipate excess light energy. Poorer PSII performance led to these responses during transitions between high and low light phases. These findings suggest a hypothesis that Cyt *b_6_/f* facilitates rapid adjustments in energy dissipation pathways under dynamic light, safeguarding PSII against photodamage.

PSI performance under fluctuating light was relatively stable in the “Low” plants, with minimal changes in Y(I) and Y(NA). However, the increases in Y(ND) reflect large donor-side limitations on the PSI electron flow (**Figure 4**), resulting from reduced electron flow through Cyt *b_6_/f*. These results underscore the robustness of PSI under fluctuating light. The maintenance of PSI function despite the severe reduction in Cyt *b_6_*/*f* content may be due to a greater contribution from alternative electron pathways.

### A Novel Trade-Off Between Photosynthetic Capacities and Resistance to PSI Photoinhibition

One of the most unexpected findings was the exceptional resistance of PSI in “Low” plants to photoinhibition under extreme fluctuating light conditions. While WT plants experienced significant declines in P*_m_* at high light intensities, the “Low” plants maintained stable P*_m_*, even at 20,000 μmol m⁻² s⁻¹ (**Figure 5**). This suggests that the reduced Cyt *b_6_/f* content alters the balance of electron flow, leading to conditions that favor PSI stability.

In our previous work, sunflower (*Helianthus annuus*) has showed a marked tolerance of PSI to fluctuating light with 800 ms-high light pulses at 5000 µmol m⁻² s⁻¹ (Kono et al. 2017). This sunflower’s ability to withstand high fluctuating light may be explained by its high-light acclimation and robust photoprotection mechanisms (Schottler and Toth 2014; Matsubara et al. 2016; Alboresi et al. 2018). The tolerance of sunflower PSI under fluctuating light reflects the “Low” transgenic tobacco plants in their PSI photoprotection but contrasts in photosynthetic capacity due to higher electron transport rate and better NPQ regulation in sunflower. The specific reduction in the Cyt *b_6_/f* complex content in the tobacco plants indicates that, while PSI tolerance was improved, overall photosynthetic capacity was weakned. Sunflowers may circumvent this trade-off through their naturally higher abundance of the Cyt *b_6_/f* complex and more efficient photoprotective mechanisms

In contrast, PSII in both WT and “Low” plants exhibited similar photoinhibition under extreme light intensities. This finding indicates that Cyt *b_6_/f* plays a more selective role in protecting PSI under high light stress. This trade-off between reduced photosynthetic capacity and enhanced PSI robustness provides a potential avenue for improving crop tolerance to extreme light fluctuations, particularly in environments prone to high irradiance.

### The Physiological Significance of Cyt *b*_6_*/f* in Dynamic Environmental Conditions

Fluctuating light is a feature of natural environments, where plants are exposed to dynamic irradiance caused by sunflecks and cloud cover (Pearcy 1983; Chazdon 1988). Our findings offer several considerations, particularly in terms of addressing the physiological significance of Cyt *b_6_/f* under natural light conditions (specially, see below). This is particularly important for understanding photosynthetic performance in crops growing under heterogeneous light environments (Vierling and Wessman 2000; Leakey et al. 2003).

In constant weak light environment, photosynthetic electron transport and photoprotection mechanisms operate under relatively low stress, as there is minimal risk of photoinhibition, and thus the ability to maintain steady-state photosynthesis is critical. The modest demand for electron transport aligns with the limited availability of light energy, minimizing stress to PSI and PSII. Plants with moderately reduced Cyt *b_6_/f* content may perform adequately, though their growth potential may be constrained.

In constant high light, the risk of photoinhibition is significantly increased due to the sustained energy input exceeding the electron transport capacity (Demmig-Adams et al. 1996; Kono et al. 2022). WT plants or those with normal Cyt *b_6_/f* levels excel in such environments, as Cyt *b_6_/f* ensures efficient electron transport and energy dissipation through robust *pmf* generation and NPQ activation. Increased NPQ in WT plants protects PSII under prolonged high light, reducing photodamage. In contrast, as shown in the present study, “Low” plants exhibit poor NPQ induction and weaker regulation of PSII and PSI activity (**Figures 2, 3, S1 and S2**), making them vulnerable to photodamage. Therefore, WT plants with optimal Cyt *b_6_/f* content are better suited to constant high light environments. Protection capability of “Low” plants is limited to PSI, making them less competitive in such conditions.

Fluctuating light dominant on the deep forest floor is characterized by sunflecks (strong light lasting for milliseconds or seconds) superimposed on a background of weak light for extended periods (Pearcy 1990; Durand et al. 2021). In such an environment, the challenge is to quickly adapt to the temporary strong light, that is, to avoid photoinhibition and maximize the use of light for photosynthesis (Kono and Terashima 2014). PSI photoinhibition resistance observed in “Low” plants is a significant advantage in this environment. The transient nature of high light ensures that reduced Cyt *b_6_/f* content does not severely limit overall photosynthesis, as PSI stability is preserved. Reduced ΔΨ in “Low” plants may favor cyclic electron flow, which could support ATP synthesis during short periods of strong light (Yamori and Shikanai 2016). On the other hand, poor NPQ regulation in “Low” plants may lead to over-reduction of PSII and photodamage during sunflecks. The trade-off between PSI photoinhibition resistance and reduced PSII photoprotection makes “Low” plants better adapted to environments where PSI stability is critical, but WT plants are superior in managing overall photoprotection.

Gradual fluctuating light in the open sites involves slow and periodic changes between strong and weak light over several tens of minutes (Valladares 2003; Miyashita et al. 2012). Plants need to balance energy use capacity and photoprotection across prolonged high and low light phases (Kono et al. 2020). WT plants with high Cyt *b_6_/f* content perform well in such environments due to their robust electron transport capacity and ability to regulate NPQ efficiently during transitions. Reduced Cyt *b_6_/f* in “Low” plants decrases PSII performance under high light and limits the capacity to dissipate excess energy effectively, as shown by delayed NPQ induction and suboptimal qL regulation.

### Broader Implications and Future Directions

The ecological significance of Cyt *b_6_/f* modulation has been demonstrated in previous studies, showing that plants adjust its content to optimize photosynthetic performance under specific environmental conditions. Terashima et al. (2021) found that the shade-tolerant plant *Alocasia odora* resists PSI photoinhibition under fluctuating light by reducing the Cyt *b_6_/f* per P700 ratio, thereby limiting electron flow into PSI. This mechanism helps maintain P700*^+^* and minimizes photodamage during transient high light. Additionally, *A. odora* plants acclimated to different light environments—constant low- or high-light and fluctuating light—exhibited varying levels of most photosynthetic components, yet the P700/chlorophyll ratio remained relatively unchanged. This suggests that maintaining a stable P700/chlorophyll ratio is crucial for PSI robustness under dynamic light conditions.

A recent paper of Terashima et al. (2025) suggests that excitation energy spillover from PSII to PSI plays a significant role in protecting both photosystems under fluctuating light conditions. Even when NPQ levels are low, excess excitation energy from PSII can be transferred to PSI, helping to maintain an oxidized P700 (P700⁺), which is crucial for preventing PSI photoinhibition. This mechanism not only stabilizes PSI by reducing over-reduction of its acceptor side but also alleviates excessive excitation pressure on PSII, thereby avoiding photodamage. The efficiency of spillover makes it a particularly advantageous photoprotective strategy compared to NPQ, which requires several minutes for activation.

The importance of optimizing photosynthetic capacity and stress tolerance in crops would meet the challenges of climate change. Our findings contribute to this goal by identifying Cyt *b_6_/f* as a key player in balancing energy transduction, photoprotection, and stress resilience. Future research should focus on unraveling the molecular mechanisms underlying PSI stabilization in “Low” plants, particularly the potential involvement of alternative electron pathways such as cyclic electron flow or chlororespiration (Nawrocki et al. 2019) etc. Investigating the interplay between Cyt *b_6_/f* and other photoprotective mechanisms, including state transitions (Shang et al. 2023), redox regulation system in chloroplast (Yoshida and Hisabori 2023), and spillover (Terashima et al. 2021, 2025; Nanda et al. 2025) could provide deeper insights into the dynamic regulation of photosynthesis. Additionally, it will be valuable to explore how the observed trade-offs between photosynthetic capacity and photoinhibition resistance manifest under field conditions. Understanding how Cyt *b_6_/f* reduction affects whole-plant performance, particularly in fluctuating natural light environments, could inform strategies for engineering crops with enhanced resilience to abiotic stress.

## Conclusions

This study highlights the pivotal role of the Cyt *b_6_/f* complex in balancing photosynthetic capacity and photoprotection under dynamic light conditions. By employing transgenic tobacco plants with reduced Cyt *b_6_/f* contents, we demonstrated that this complex is essential for maintaining the proton motive force (*pmf*) and ensuring efficient energy conversion. While moderate reductions in Cyt *b_6_/f* had limited impact on photosynthetic performance, severe reductions decreased electron transport rate and NPQ under fluctuating light. Remarkably, plants with reduced Cyt *b_6_/f* exhibited exceptional resistance to PSI photoinhibition, revealing a trade-off between maximizing photosynthetic capacity and enhancing stress resilience. These findings highlight the potential of fine-tuning Cyt *b_6_/f* levels as a strategy to improve crop productivity and tolerance to fluctuating light environments.

## Abbreviations

Cyt *b_6_/f*: cytochrome *b_6_/f*
F*_m_*: maximum fluorescence in the dark
F*_m_*’: maximum fluorescence in actinic light
F*_o_*: minimum fluorescence in the dark
F*_o_*’: minimal fluorescence yield in actinic light
Fs’: steady-state chlorophyll fluorescence level in actinic light
F*_v_*: variable fluorescence (F*_m_* – F*_o_*)
ML: measuring light
NPQ: non-photochemical quenching of chlorophyll
PAM: pulse-amplitude modulation
P700: PSI reaction centre
PPFD: photosynthetic photon flux density
PSI: photosystem I
PSII: photosystem II
P*_m_*: maximal P700 signal upon full oxidation
P*_m_*’: maximal P700 signal in the presence of actinic light
P*_o_*: complete reduction level of P700 signal
P: oxidized state P700 signal in the presence of actinic light
qL: fraction of open PSII centers
SP: saturation pulse
Y(II): quantum yield of the PSII photochemistry
Y(I): quantum yield of the PSI photochemistry
Y(ND): quantum yield of non-photochemical energy dissipation due to the donor-side limitation on PSI electron flow
Y(NA): quantum yield of non-photochemical energy dissipation due to the acceptor-side limitation on PSI electron flow

## Author contributions

M.K., I.T. and W.Y. conceived and designed the study. M.K., H.K., K.T. and W.Y. conducted experiments and performed data analysis. M.K. and I.T. and W.Y. wrote the manuscript. All authors discussed the results and revised the manuscript.

## Availability of data

Supporting data can be requested by contacting the corresponding author.

