## Supplemental Figures for "Cyt *b_6_/f* Complex Fine-Tunes PSI Stability and Photosynthetic Capacity Under Fluctuating Light"

**Fig S1.** Results of Student's t test ($P < 0.05$) for the data of Figure 2. The values represent the mean ± SE. Different letters indicate significant differences among the Rieske FeS content.

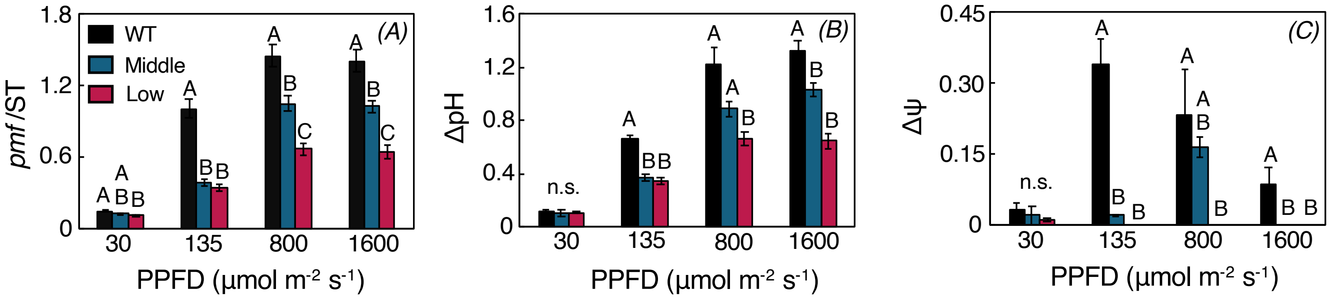


**Fig S2.** Results of Student's t test ($P < 0.05$) for the data of Figure 3. The values represent the mean ± SE. Different letters indicate significant differences among the Rieske FeS content.

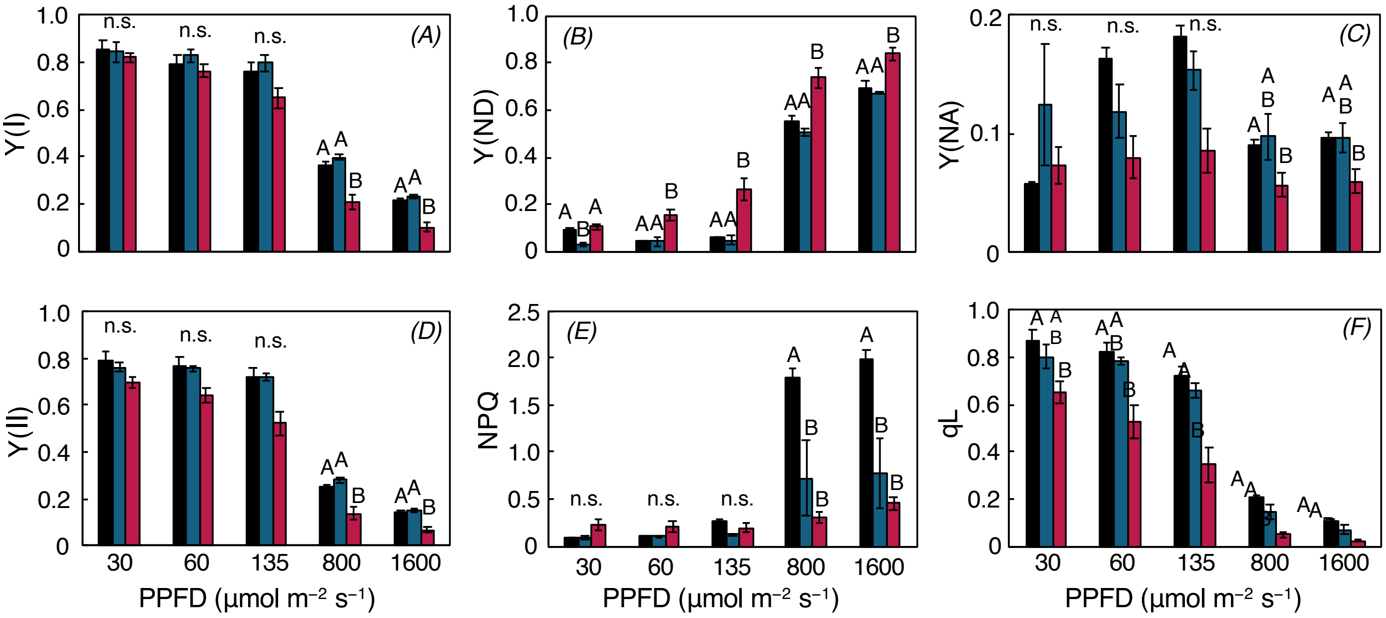
